# Recent global changes have decoupled species richness from specialization patterns in North American birds

**DOI:** 10.1101/577841

**Authors:** Anne Mimet, Robert Buitenwerf, Brody Sandel, Jens-Christian Svenning, Signe Normand

## Abstract

**Aim:** Theory suggests that increasing productivity and climate stability toward the tropics can explain the latitudinal richness gradient by favouring specialization. A positive relationship between species richness and specialization should thus emerge as a fundamental biogeographic pattern. However, land use and climate change disproportionally increase the local extirpation risk for specialists, potentially impacting this pattern. Here, we empirically quantify the richness-specialization prediction and test how 50 years of climate and land use change has affected the richness-specialization relationship.

**Location**

USA

**Time period**

1966-2015

**Major taxa studied**

Birds

**Methods:** We used the North American breeding bird survey to quantify bird community richness and specialization to habitat and climate. We assess i) temporal change in the slope of the richness-specialization relationship, using a Generalized Mixed Model; ii) temporal change in spatial covariation of richness and specialization as driven by local environmental conditions, using Generalized Additive Models; and iii) land use, climate and topographic drivers of the spatio-temporal changes in the relationship, using a multivariate method.

**Results:** We found evidence for a positive richness-specialization relationship in bird communities. However, the slope of the relationship declined strongly over time. Richness spatially covaried with specialization following a unimodal pattern. The peak of the unimodal pattern shifted toward less specialized communities over time. These temporal changes were associated with precipitation change, decreasing temperature stability and land use.

**Main conclusions:** Recent climate and land use changes induced two antagonist types of community responses. In human-dominated areas, the decoupling of richness and specialization drove a general biotic homogenization trend. In human-preserved areas under increasing climate harshness, specialization increased while richness decreased in a “specialization” trend. Our results offer new support for specialization as a key driver of macroecological diversity patterns, and show that global changes are erasing this fundamental macroecological pattern.

**Biosketch:** Anne Mimet is a postdoctoral researcher interested in the understanding of human impacts on biodiversity through land use and climate changes, at various spatio-temporal scales. She is interested in embracing the complexity of socio-ecological systems, and in the understanding of biodiversity trends in a human-dominated world in the context of the general theories of ecology.

## Introduction

Climate is a major driver of global diversity patterns (Araújo et al., 2008; Dynesius & Jansson, 2000; Mannion, Upchurch, Benson, & Goswami, 2014). In the context of the global biodiversity crisis, understanding how human-induced land use and climate changes have and will change macro-ecological patterns is crucial in planning biodiversity and ecosystem conservation (Flojgaard, Normand, Skov, & Svenning, 2011; Sandom, Faurby, Sandel, & Svenning, 2014).

Numerous hypotheses have been proposed to explain the latitudinal gradient of species richness (Willig, Kaufman, & Stevens, 2003). These hypotheses can be classified into two nonexclusive categories, either emphasize the importance of ecological or evolutionary processes in driving the pattern. On one hand, the ecological hypotheses propose that the latitudinal richness gradient is driven by gradients of niche-breadth, species interactions, range size, or area. On the other hand, the historical hypotheses propose that the latitudinal richness gradient is driven by diversification rates, and some historical hypotheses explain the latitudinal richness gradient by time for diversification and colonization (Currie et al., 2004; Gaston, 2000; MacArthur, 1972; Stevens, 1989; Svenning & Skov, 2007).

Regardless of the primary driving force, most theories predict that specialization increases towards the species-rich tropics due to higher diversification rates, caused by either higher productivity and/or long-term environmental stability (Belmaker, Sekercioglu, & Jetz, 2012; Currie et al., 2004; Dynesius & Jansson, 2000; MacArthur, 1972). Hence, the hypotheses generally predict a positive relationship between richness and specialization. However, the richness-specialization relationship in itself has only rarely been studied, finding support in some studies (Belmaker et al., 2012; Dalsgaard et al., 2011), but no evidence from another (Novotny, 2006). Support for a positive richness-specialization relationship is mainly based on indirect evidence from studies that explore the latitude-specialization and latitude-range size gradients. These indirect studies use latitude as a proxy of richness or as a proxy for the underlying drivers of productivity and long-term stability, and their results does not always support a latitudinal specialization gradient (Beaver, 1979; Dalsgaard et al., 2011; Novotny, 2006; Schleuning et al., 2012). Results from these direct and indirect approaches paint an unclear picture of the reality of the richness-specialization relationship. These opposing results highlight the need to quantify the relationship itself rather than using latitudinal proxies for richness, productivity or long-term stability (Gaston, 2000).

Human-driven global changes may modify crucial ecological processes and macroecological patterns through changes in available productivity, environmental stability and heterogeneity. For example, land use and climate changes strongly impact available productivity (Haberl et al., 2007; Melillo et al., 1993; Newbold et al., 2018) and environmental stability through net changes in biotic and abiotic conditions (e.g. mean temperature, precipitation and available productivity), but also through increases in inter-annual variability in these conditions (Chamberlain, Fuller, Bunce, Duckworth, & Shrubb, 2000; Jentsch, Kreyling, & Beierkuhnlein, 2007). Furthermore, land use and climate changes modify spatial heterogeneity in … conditions and thus available habitat, which is positively correlated to richness (Allouche, Kalyuzhny, Moreno-Rueda, Pizarro, & Kadmon, 2012; Stein, Gerstner, & Kreft, 2014).

Perhaps counter-intuitively, global changes have not systematically led to local reductions in species richness over the last decades, and trends in species richness are often not good measures of changes in biodiversity (Hillebrand et al., 2017). The inconsistent response of species richness to land use changes in different contexts can be partly explained by a trade-off between changes in the proportion of used land and changes in landscape heterogeneity, which have opposite impacts on local species richness (Allouche et al., 2012; Pellissier, Mimet, Fontaine, Svenning, & Couvet, 2017; Stein et al., 2014). Instead of consistently decreasing richness, land use and climate changes tend to increase communities’ turnover (Hillebrand et al., 2017), typically governed by the spread of winner species (ruderal, generalist) and the loss of late successional, specialist and endemic species (Allouche et al., 2012; Devictor et al., 2008). Together, these trends lead to worldwide biotic homogenization (Davey, Chamberlain, Newson, Noble, & Johnston, 2012; McKinney & Lockwood, 1999).

In this paper, we aim to validate the existence of the richness-specialization relationship for bird communities in North America and test the hypothesis that the relationship has weakened in the last decades because of land use and climate changes, leading to biotic homogenization. We also aim to assess if the relationship can be present, but hidden by other spatial patterns, by contrasting local conditions of variables linked to productivity and stability. Productivity and long-term environmental stability are indeed independently driving species richness and specialization. The positive relationship between primary productivity and species richness, the so-called species-energy relationship, has found considerable support for many taxa from local to global scales (Gillman et al., 2015; Wright, 1983). Productivity is also identified as a driver of specialization (Abrams, 1995; Pellissier, Barnagaud, Kissling, Sekercioglu, & Svenning, 2018). Environmental stability is required for species specialization to emerge (Dynesius & Jansson, 2000; Ponge, 2013; Stevens, 1989) and be maintained (Allouche et al., 2012; Devictor et al., 2008; Ponge, 2013). Spatial variations in local conditions of productivity and environmental stability are thus expected to drive important changes in species richness and specialization that are likely to hide the richness-specialization relationship. In other words, the shape of the spatial covariation of productivity and environmental stability should be the main driver of the spatial covariation of species richness and specialization. A non-linear spatial covariation of productivity and environmental stability should therefore lead to a non-linear spatial covariation of richness and specialization. We expect to observe such a non-linear spatial covariation pattern for birds communities in the United States, with low richness and low specialization in highly homogeneous anthropogenic landscapes; high species richness but low specialization in heterogeneous and productive landscapes under intermediate human pressure; and high specialization but low richness in harsh environments little impacted by human activities, such as many mountainous areas. We expect to see richness and specialization co-vary in space due to variables that regulate the quantity and diversity of resource and modify environmental stability, and not necessarily to follow a latitudinal gradient.

We used 50 years of data from the North-American Breeding Bird Survey (1966-2015) (Figure 1). We first model the change over time of the richness-specialization relationship removing the effect of local conditions on richness. We then explore the shape of the spatial covariation of richness and specialization, as driven by local conditions, and how this shape has changed in the past 50 decades. We finally identify the past and present land use, climate and topographic conditions that explain the spatial and temporal changes in the richness-specialization relationship using a multivariate method adapted to spatio-temporal analyses.

**Figure 1:**
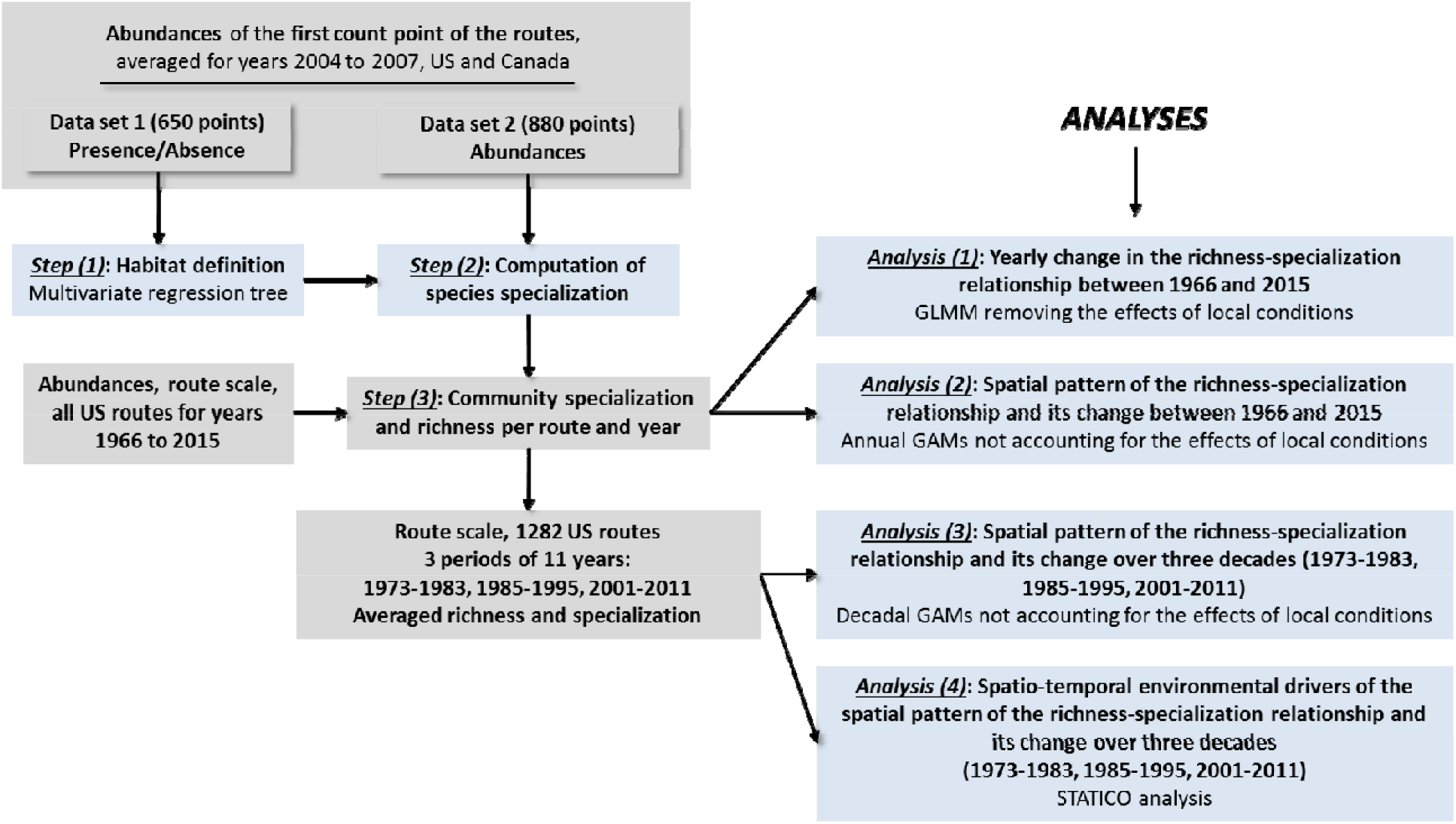
Workflow implemented in the study. The data are represented in grey and computation/analyses are represented in light blue. The steps and analyses numbers refer to the same numbers in the method and results sections.

## 2. METHODS

### 2.1 Bird data

We quantified bird species richness and habitat specialization based on the North American Breeding Bird Survey (BBS), which is a database of standardized breeding bird abundances covering 1966 to 2015 (see Appendix S1 for more information). The standard observational units of the BBS are routes of approximately 42 km long, composed of 50 count points each, where only the first count point has specified geographic coordinates. Routes are surveyed once a year, but not all routes are surveyed all years. The number of surveyed routes increases through time, from 533 in 1966 to 2760 in 2015.

We used different subsets of the route data for different parts of this study (Figure 1): the first count point of some routes (2004 to 2007) was used to compute species-level specialization; the yearly route data was used to study the changes of the richness-specialization relationship over years; and the route data was averaged over decades (1973-1983,1985-1995, 2001-2011) to study the change of the relationship at the decadal scale and identify the environmental variables linked to those changes (Figure 2).

**Figure 2:**
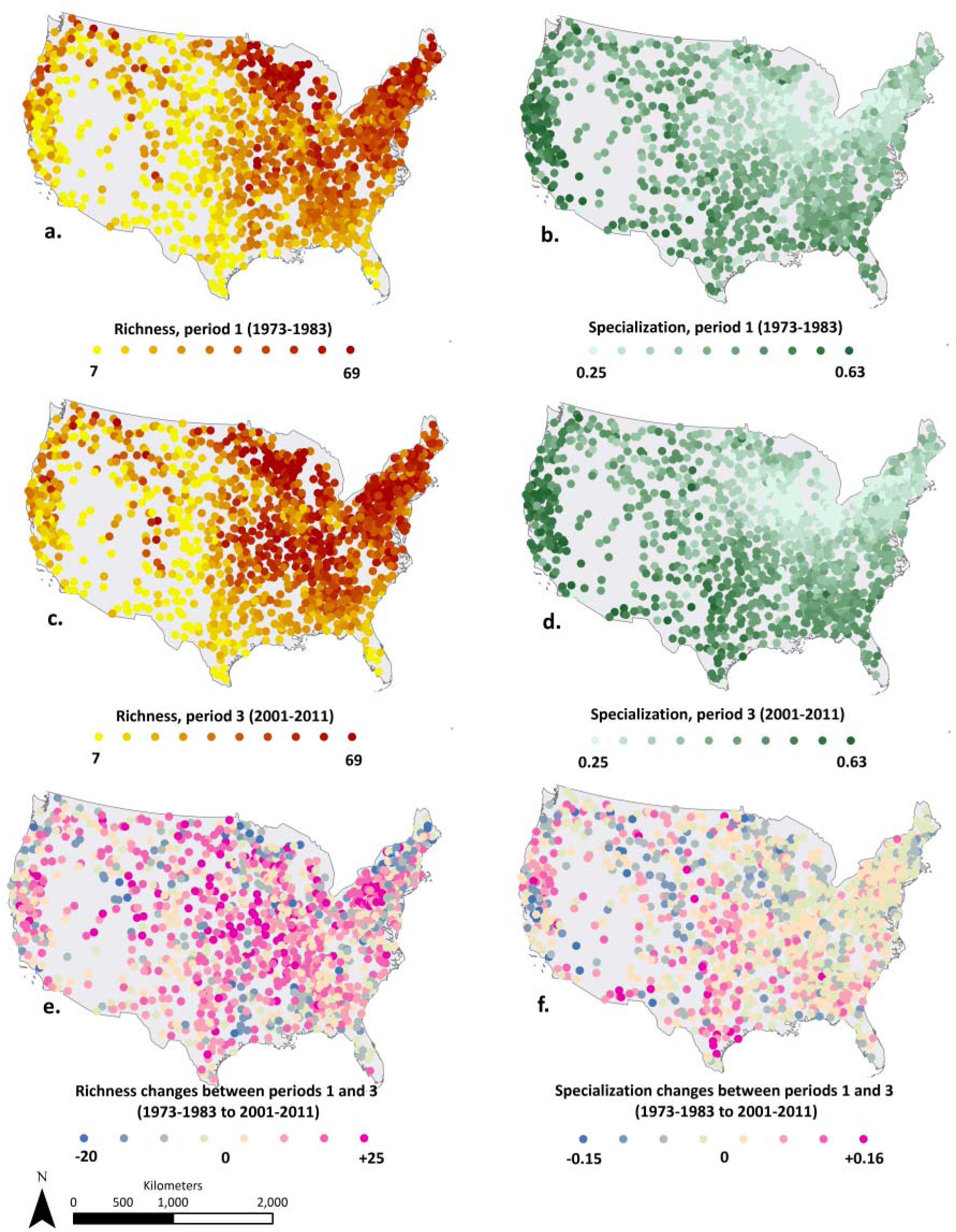
Richness and specialization of breeding birds in the first decade (1973-1983) and third decade (2001-2011) and the changes in richness and specialization between the first and the third decades.

### 2.2 Species and communities specialization to climate and habitat

Species-level specialization (niche breadth) was estimated accounting for both climate and habitat. We computed the community-level specialization following a three-step approach (Figure 2). Step (1): We defined ecologically meaningful habitat types based on community response to land cover and climate; step (2): we computed species-level specialization based on the species abundances within these habitat types; step (3): we quantified community specialization as the abundance-weighted mean of species-specific specialization values.

Since only the first point of each route has geographic coordinates, we used the first count points of BBS data routes for the years 2004 to 2007 to obtain species-level specialization (steps (1) and (2). We included Canadian data, but excluded count points in urban areas and wetlands, for which the number of observations was too low to obtain relevant specialization estimates (see Appendix xx for data preparation). We extracted the land cover and climate of each count point from the corresponding pixels respectively in the GlobCover land cover product (2006, 300 m resolution; http://due.esrin.esa.int/page_globcover.php) (simplified nomenclature, Appendix S2) and the Köppen climate classes (Peel, Finlayson, & McMahon, 2007). The simplification of the land cover nomenclature resulted in seven land cover classes, providing comparable ecological grain to short (grasslands, croplands) and tall vegetation (different forest types) land covers (Devictor et al., 2008): croplands (138 count points), herbaceous vegetation (395 count points), sparse vegetation (59 count points), shrubland (168 count points), broadleaved forest (269 count points), needleleaved forest (380 count points) and mixed forest (121 count points). We restricted the analyses to Köppen climate zones containing more than 15 count points, which included11 climate zones (Figure 3).

**Figure 3:**
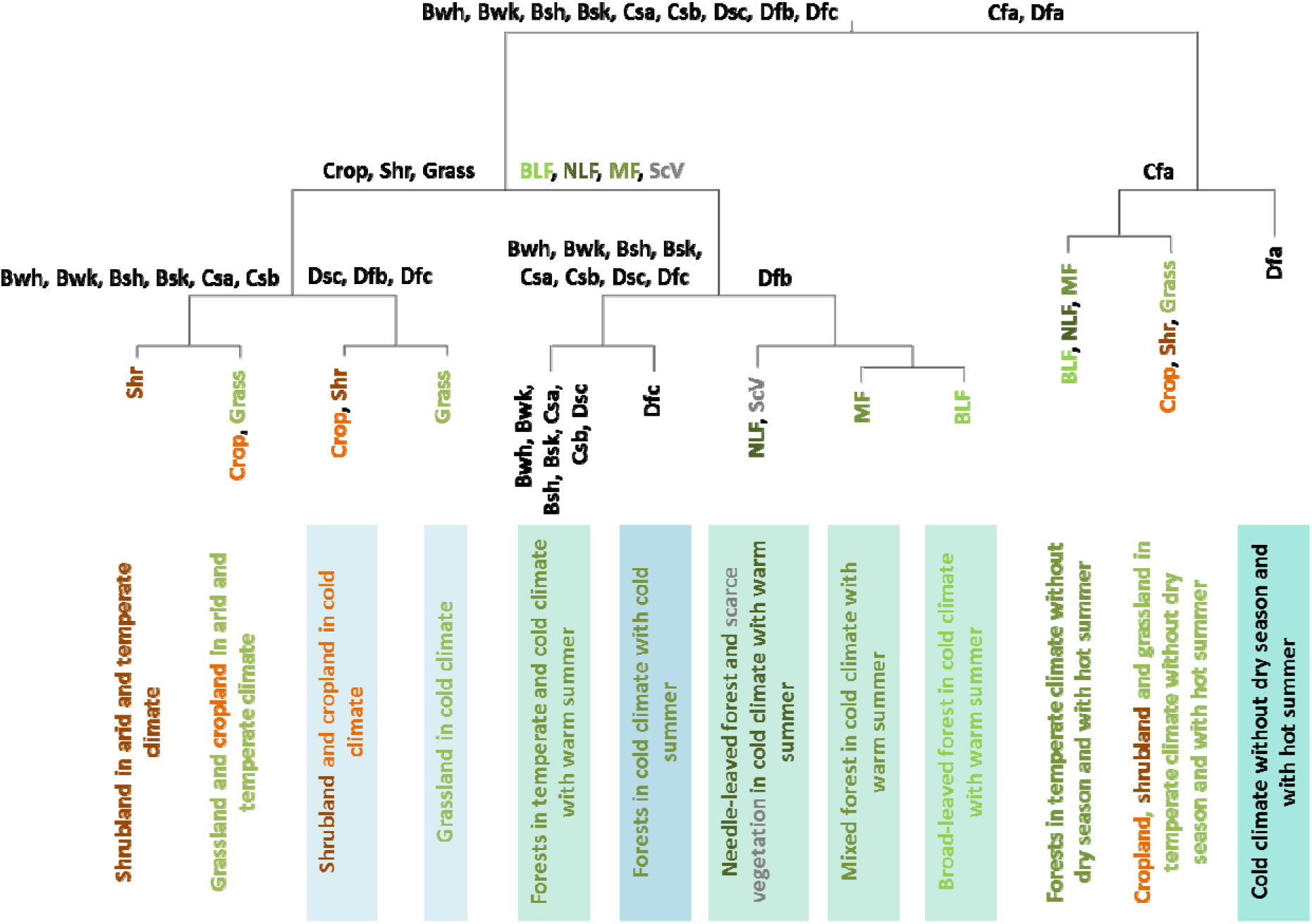
Habitat types obtained from land cover and Köppen climate classes. The Köppen climate zones appear in black and land covers in colors. Köppen climate zones correspond to: Bwh: Arid hot desert; Bwk: Arid cold desert; Bsh: Arid hot steppe; Bsk: Arid cold steppe; Csa: Temperate, dry ad hot summer; Csb: Temperate, dry and warm summer; Cfa: Temperate, without dry season, hot summer; Dsc: Cold, cold and dry summer; Dfa: Cold, without dry season, hot summer; Dfb: Cold, without dry season, warm summer; Dfc: Cold, without dry season, cold summer. Land cover classes correspond to: Shr: shrubland; Crop: cropland; Grass: grassland; NLF: Needle-leaved forest; ScV: Scarce vegetation; MF: Mixed forest; BLF: Broadleaved forest. The boxes at the bottom summarize the climate and land cover information of the habitats created by the tree.

We took several actions on data selection to achieve steps (1) and (2) avoiding circularity issues linked to the computation of the species-level specialization. First, we only used for steps (1) and (2) count points that were not used in the analyses explaining community-level specialization by environmental variables, leading to a selection of 1530 count points. Second, to avoid using the same data to define the habitat types (step (1)) and the species-level specialization (step (2)), we randomly split those 1530 count points into two independent datasets of 650 and 880 count points. We kept more data for step (2) in order to obtain better estimations of species specialization, while after several tests the structure of the tree appeared robust to reduced data input.

#### 2.2.1. Step (1): define ecologically meaningful habitat classes for bird communities

We used a multivariate regression tree to define ecologically meaningful habitat types for bird communities. The tree identified combinations of land cover and climate classes that best explained observed species presence/absence (mvpart package in R) (Therneau & Atkinson, 2013). For defining ecologically meaningful habitat classes we only considered species present in more than five count points (153 species).

To avoid overfitting, we pruned the tree using cross-validation (CV Error) (Borcard, Gillet, Legendre, & Legendre, 2011) and retained 11 splits that led to 12 leaves (Figure 3). Each leaf was characterized by at least one land cover and climate class, with the exception of one class, which was defined solely based on climate (cold climate with warm summer).

#### 2.2.2. Step (2): Computing species-level specialization

Species-level specialization was computed on a set of 880 count points independent of the set used for defining the habitat classes (step 1). We only computed specialization for species recorded at a minimum of five count points, which resulted in species-level specialization estimates for 162 bird species.

Species affinity to each habitat was computed as the frequency of occurrence of a species in each habitat typeas defined in step (1). Affinity is described as the indicator value A in Dufrene & Legendre (1997) (Eq 1).

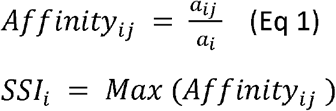

where a_ij_. is the abundance of individuals of species *i* observed in the habitat type *j* and a_i_ is the total abundance of species *i* across all sites.

The affinity of a species to a habitat type is constrained between 0 (the species is never observed in a certain habitat type) to 1 (the species in exclusively observed in a certain habitat type). Species-level specialization (SSI) was then defined as the maximum habitat affinity across all 12 habitat types (Equation 2)(Figure 3; Appendix S4).

#### 2.2.3. Step (3): Computing community-level specialization and richness

Community-level specialization was computed at the route level, i.e. considering the total abundance of the species observed over the 50 count points of each route. A community-level specialization value was computed for all routes sampled at least once between 1966 and 2015 in the United States only. Following previous studies, we defined community-level specialization (CSI) as the community-weighted mean species-level specialization score (Devictor et al., 2008) (Eq 3).

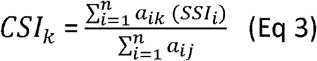

Where *CSI_k_* is the specialization of community *k*, n is the total number of species, *a_ik_* is the abundance of individuals of species *i* in community *k*, and *SSI_i_* is the specialization of species *i*. We computed the CSI for each route and each year of the study. Species richness was simply defined as the total number of species observed in a route per year.

### 2.3. Spatiotemporal changes in the richness-specialization relationship

For the following analyses, we used the BBS data of the US at the route scale, for the years 1966 to 2015. The richness-specialization relationship was thus computed for bird communities, the term “richness” referring to species richness and the term “specialization” referring to the community-level specialization.

#### 2.3.1. Analysis (1): Temporal change in the richness-specialization relationship

We quantified the richness-specialization relationship and its temporal change using a Generalized Mixed Model (GLMM), where species richness was explained by year (defined as a categorical variable), community-level specialization and their interaction (Equation 3). The interaction allows us to estimate how the slope of the richness-specialization relationship varies by year. Route was specified as the random factor to account for the effects of local conditions on the estimation of the relationship. We residual spatial autocorrelation was non-significant for most years, and under 2% for the others, suggesting that the predictor variables sufficiently captured any spatial dependence in the response variable (spline.correlog, ncf package) (Bjornstad, 2016).

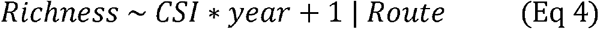

#### 2.3.2. Analyses (2) and (3): Spatial covariation of the richness and specialization and its temporal change

Local conditions linked to productivity and environmental stability are the very important drivers of richness and specialization, and the way they covary in space is likely to visually hide the richness-specialization relationship and induce a very different spatial covariation pattern of the relationship. To assess how local conditions shape the covariation of richness and specialization relationship in space, we explored the spatial patterns of the richness-specialization relationship and their changes over time using spatial models that did not account for local conditions.

To do so, we used Generalized Additive Models (GAMs) to predict richness using specialization at two temporal scales: annual and decadal (11 years). We defined the three decades based on available land use from the HYDE data base (Appendix xx): 1973-1983,1985-1995 and 20012011. For the decadal analysis we only selected routes that were sampled at least three times within each decade, which resulted in a data set of 1282 routes. Within each decade, species richness and community-level specialization estimates were averaged for each route. To identify the shape of the covariation of richness and specialization, we fit linear and piece-wise models with one and two breakpoints. For the piecewise regressions, we identified the best breakpoint locations (and the best combination of breakpoints for the two-breakpoints models) using an iterative approach that searches for the model that maximised explained deviance (Crawley, 2007). Once all models were fitted, we selected the best model for each year/decade using AIC values.

Spatial autocorrelation, i.e. how the value a variable at a location varies together with its value at other locations, depends in ecology on three main causes (Dormann et al., 2009): 1) ecological processes are distance-dependent, 2) the shape of the relationship between environment and species is not modelled correctly, and 3) important covariates are missing from the statistical model. In the present set of analyses, we excluded in purpose the covariates (variables linked to productivity and environmental stability) from the models to look at the shape of the richness-specialization relationship as driven by those important variables. It is thus expecting to observe residual autocorrelation in the models. However, the other causes of spatial autocorrelation still need to be accounted for in the model. We used a GAM approach which simultaneously allowed us to look for the appropriate shape of the covariation pattern by comparing linear and piecewise regression models (see above) (cause 2) and to explicitly represent spatial autocorrelation linked to richness values at nearby locations by including a spline interaction term of the latitude and longitude (cause 1). The temporal regressions in slopes were inversely weighted by the squared standard deviation of the slopes annual values.

We quantified temporal trends in the model parameters (i.e. slopes, locations of the breakpoints and distance between the breakpoints) using linear regressions.

### 2.4. Analysis (4): Spatio-temporal environmental drivers of spatial pattern of the richness-specialization relationship

We attributed spatio-temporal changes of the richness-specialization relationship between 1973 and 2011 to environmental drivers. In this analysis we used abundance data within the three decades (1973-1983,1985-1995 and 2001-2011).

#### 2.4.1. Environmental drivers

Four types of environmental variables were explored. All variables were computed in a 9500m buffer around the routes (buffer size based on the resolution of the HYDE land cover data, Appendix S5). (i) Land use: the proportion of each land use within the buffer area, naturalness, irrigation and human population density (HYDE) (Klein Goldewijk, Beusen, Van Drecht, & De Vos, 2011), (ii) climate, capturing the multiple dimensions of climate change (Garcia, Cabeza, Rahbek, & Araújo, 2014; Waldock, Dornelas, & Bates, 2018): mean, minimum, and spatial and temporal range of the minimum and average temperature and precipitation during winter (January and February) and breeding season (May and June) (PRISM; http://prism.oregonstate.edu/), (iii) topography: elevation and the standard deviation of elevation (SRTM30+, http://glcf.umd.edu/data/srtm/), (iv) long-term Net Primary Productivity (NPP0) (Haberl et al., 2007), and (v) climate stability; measured as climate velocity during the Pleistocene and obtained from (Sandel et al., 2011).

#### 2.4.2. Multivariate analysis of drivers of change

We attributed changes in the richness-specialization relationship to environmental drivers using a STATICO analysis. This multivariate method makes possible to analyse a series of paired ecological tables through time. The STATICO analysis identifies the common structure between the response and the environmental variables tables through time, termed the common space (see Appendix S6 for a comprehensive description of the STATICO approach). The projection of the response variables (i.e., richness and specialization) and of environmental predictors in the common space provides information about the spatial relationships between these two sets of variables. The changes in the location of the response variables in the common space reflect change of the richness-specialization relationship through time. Changes in the location of the environmental variables in the common space indicate which environmental variables are linked to temporal changes in richness and specialization.

For this analysis, we used richness and specialization in the response table, i.e. not their relationship. This allowed us to i) identify the environmental variables correlated to richness and specialization, ii) visualize the richness-specialization relationship and the variables linked to both richness and specialization in the common space, and iii) understand how temporal changes of the relationship are related to changes in richness, specialization and environmental variables. The analysis was implemented in the R package *ade4* (Simier, Blanc, Nandris, & D, 1999; Thioulouse, 2011).

## 3. RESULTS

The general temporal trends in richness and specialization, computed at the decadal scale between the first and the third decade on the 1282 routes, show that species richness increased in average from 39.7 to 42, while community-level specialization remained constant (0.35)

### 3.1. Analysis (1): Temporal changes in the richness-specialization relationship

The GLMM showed a positive relationship between richness and specialization over the entire study period. However, the slope of the relationship decreased between 1966 and 2015 (Figure 4). In other words, throughout the study period, average specialization of a community was higher in communities with more species, but specialization of a community in 2015 was lower than an equally species-rich community in 1966. This dampening effect was less strong after 1990 (Barnagaud, Gaüzère, Zuckerberg, Princé, & Svenning, 2017).

**Figure 4:**
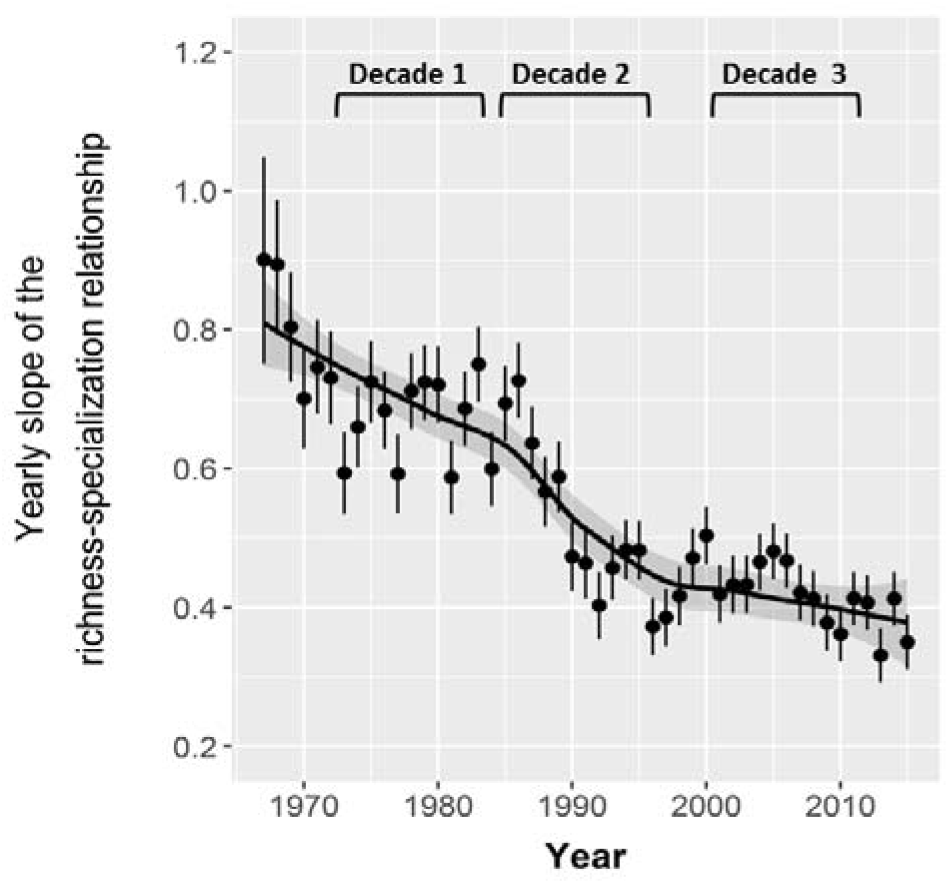
Temporal change in the slope of the richness-specialization relationship. The three decades used in some subsequent analyses are shown (1973-1983,1985-1995, 2001-2011).

### 3.2. Analysis (2) and (3): Changes over time in the spatial covariation pattern of richness and specialization

In both annual and decadal models, the spatial covariation pattern of richness and specialization was best explained by a piecewise linear regression with two breakpoints (Figure 5a; Appendix S6).

**Figure 5:**
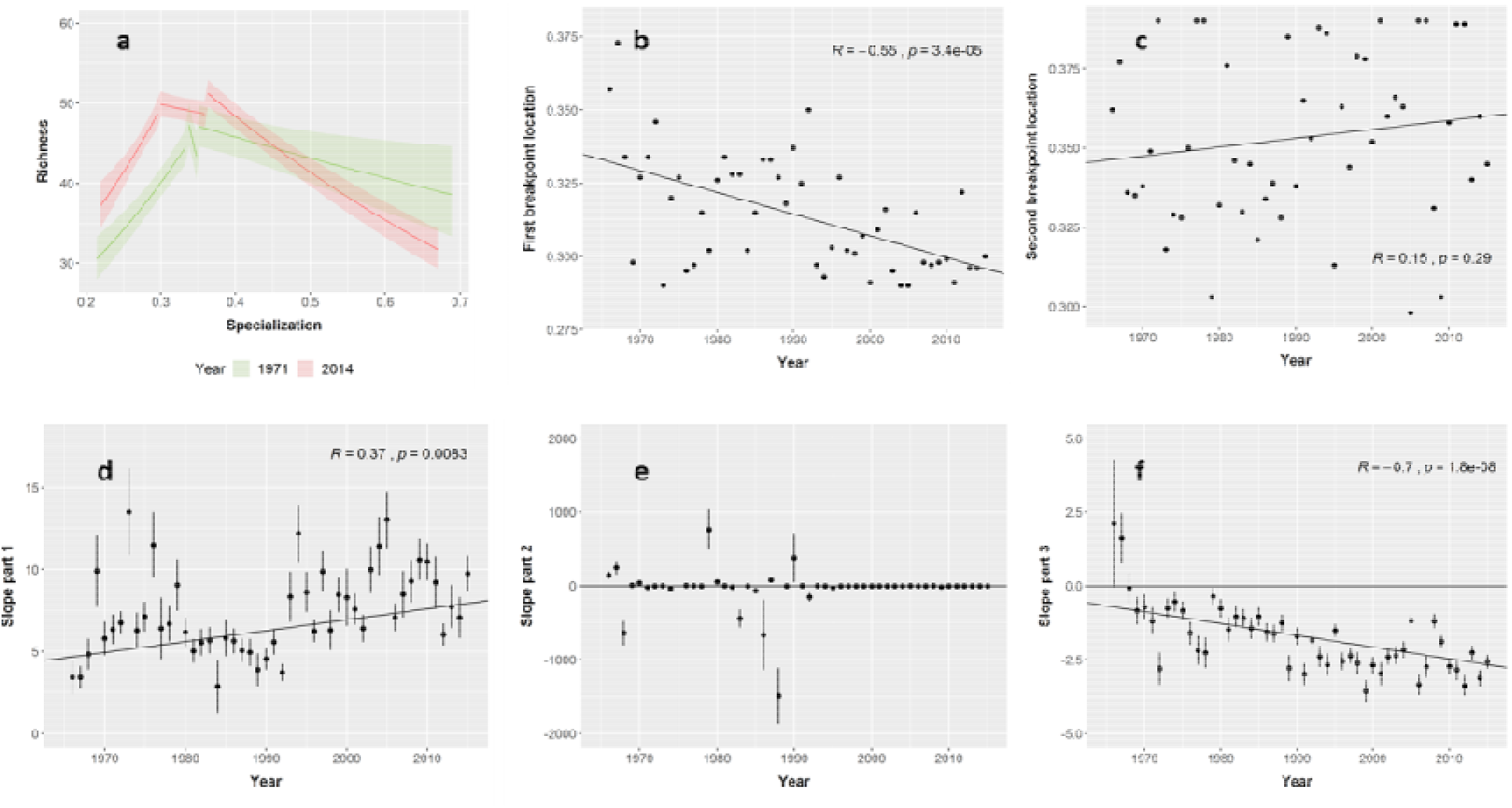
Temporal changes in the spatial shape of the richness-specialization relationship at individual routes. According to model outputs, the change is described using a two-breakpoint piecewise regression. a) Examples of modelled relationship for two years chosen to illustrate the changes in the spatial shape of the relationship between the beginning (1975) and the end (2014) of the study. We chose these two years because their breakpoints and slope values were close to the temporal regression lines, b) and c) Temporal changes in the location of the breakpoints, d), e) and f) Slopes of the three parts for the two-breakpoint piecewise regression: the error bars in the slopes figures indicate the standard errors of the coefficients. Significant temporal trends are indicated by stars.

The temporal changes in the breakpoints location and in the slopes showed how the spatial richness-specialization pattern of covariation between richness and specialization changed over time. The location of the first breakpoint significantly shifted toward lower specialization values over time (from 0.33 to 0.3), while the temporal trend of the second breakpoint was nonsignificant (Figure 5 a and b). Meanwhile, the first and third slopes significantly changed through time (Figure 5a, d, f). The first slope was positive and increased over time (Figure 5d). In contrast to the first slope, the second slope was negative and decreased over time in annual analysis but not in the decadal one. These changes in the breakpoint location and slopes cooccurred with a rise in the higher level of richness, which increased from 47 species at the beginning of the study to 51 at the end (Figure 5a).

### 3.3. Analysis (4): Environmental drivers of the spatial richness-specialization patterns and the temporal changes in the richness-specialization relationship

The ratio of common variance between the response and environmental variables on the total variance, captured by the STATICO analysis, varied from 0.26 (second decade) to 0.30 (third decade) (Monte Carlo permutation tests).

Both components of the STATICO analysis captured gradients in environmental as well as human-driven variables (Figure 6a). The first axis captures a negative correlation between richness and specialization, corresponding to the third slope in the piecewise regressions (Figure 5a). Along this axis, specialized, but species-poor communities were associated with mountainous areas, characterized by low-intensity land use and dry climates with high spatial climatic variability due to topography, as well as high long-term climate stability (i.e. low climate velocity). These communities were predominantly located in deserts and the Rocky Mountains (left half of Figure 6a; Figure 6c; Figure 7a). In contrast, species-rich and low-specialized communities were linked to the highly productive environments of the Midwest and along the East Coast, where agriculture dominates the landscape, and long-term climate instability is high (right half of Figure 6a; Figure 6c; Figure 7a). The second axis captures a gradient of richness weakly positively correlated to specialization, corresponding to the first part of the piecewise regression (Figure 5a). Along this gradient, species-rich and specialized communities are found in areas of low-intensity land use and wet climate (East Coast and Appalachian Mountains, lower half of Figure 6a and c, Figure 7b). Species-poor and low-specialized communities were associated with warm and dry areas of land intensively used for agriculture, e.g. land brought under irrigation in the South-West (upper half of Figure 6a; Figure 6c).

**Figure 6:**
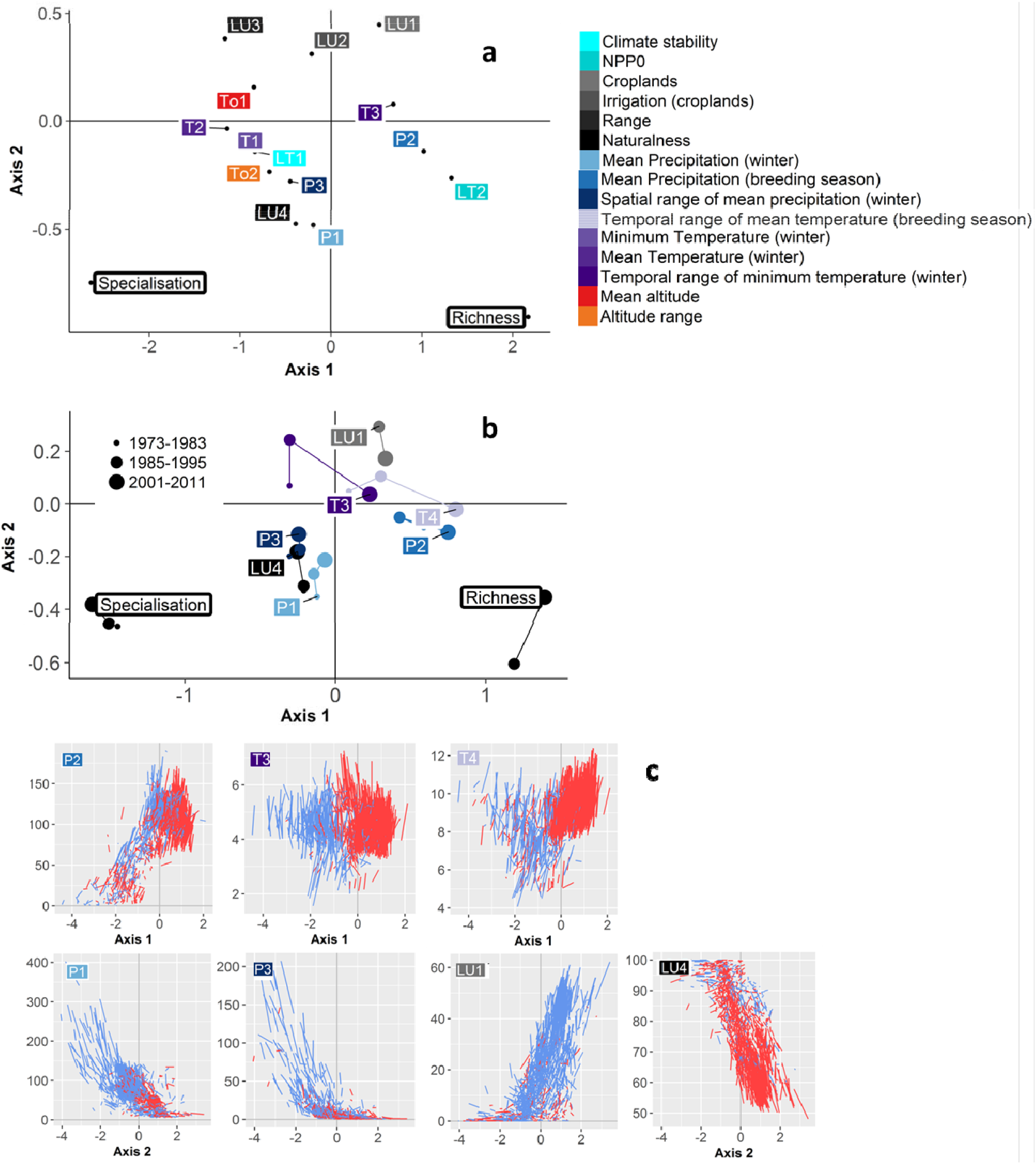
Environmental drivers of spatio-temporal change in the richness-specialization relationship. a, b) The variables are projected in the common space defined by the two axes of the STATICO analysis. Richness and specialization are shown in white boxes. Environmental variables are shown in color, the associated code in the figures stand for: **LU**: land use variable; **To**: topographic variable; **T**: Temperature related variable;; **P**: precipitation related variable; **LT:**long-term variable. **a) Association between richness, specialization and environmental variables in a twodimensional** common space. Only the environmental variables with a correlation over 20% with the components are shown and used for the interpretation of the common space. **b) Temporal dynamics of richness, specialization and environmental variables in the** common space. Small points represent the first decade (1973-1983), medium size points the second (1985-1995) and large point the third decade (2001-2011). Note that some points are hidden. Only the environmental variables with a change in correlation with one of the components over 10% are shown and used for interpreting changes in richness and specialization. **c) Temporal dynamics of the environmental variables identified in the STATICO analysis along the two axes of the STATICO analysis.** These figures represent for each route the initial value of the variable and its trend in time, making apparent how the routes are organized along the STATICO axes regarding this values and trend. We can observe very strong differences in dynamics depending on the route location on the axis, i.e. T3 and T4 along the first axis, and in initial value and trend for some others, i.e. P1 along the second axis. The dynamics are represented along the axis along which the change in correlation was observed. Red lines represent an increase in the environmental variable in time, while blue lines represent a decrease. The dynamics are computed between the first (1973-1983) and the third (2001-2011) decade.

**Figure 7:**
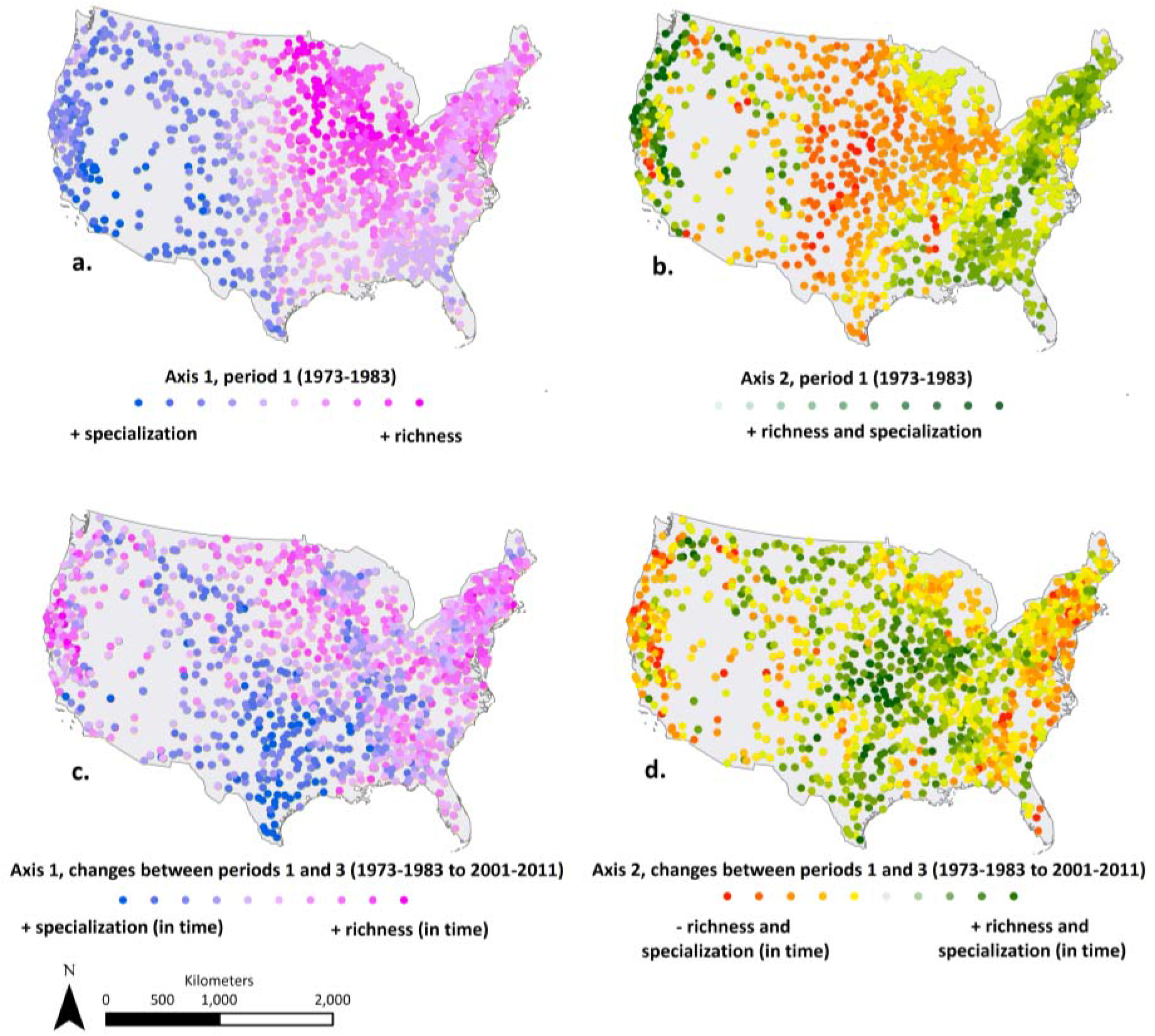
Position and temporal change of each route along the STATICO axes according to their richness and specialization: a and b. for the first decade (1973-1983); c and d for the temporal changes between the first and the third decade (1973-1983 to 2001-2011).

The shifts of richness and specialization over time (Figure 6b, Figure 7c and 7d) indicate a temporal change in the richness-specialization relationship (Appendix S7 for a more detailed explanation). The negative relationship between richness and specialization captured by the first axis (Figure 6a) became stronger in time, as richness and specialization moved away from each other (Figure 6b). Climatic stability (inter-annual variations in minimum and mean temperature) and precipitation during the breeding season strongly changed over time along the first axis. In rich, but low-specialized communities of the right part of the first axis, the increase in those variables favoured richness at the expense of specialization (Figure 6b). On the contrary, in the species-poor but specialized communities of the left part of the first axis, the decrease in those variables favoured specialization at the expense of the richness. The weakly positive relationship between richness and specialization captured by the second axis weakened over time, as land use and climate conditions slightly homogenized (Figure 6c). Thus, in the species-poor and generalist communities of warm and dry regions with intensive land use (upper half of Figure 6b; Figure 7d), this weakening was related to an increasing naturalness, which benefited both richness and specialization. In the more species-rich and specialized communities of the humid regions (lower half of Figure 6b, Figure 7d), the weakening of the relationship was related to a decrease of precipitation, negatively impacting both richness and specialization.

## DISCUSSION

### Weakening of the richness-specialization relationship through time

We find a positive richness-specialization relationship in bird communities at the continental extent, which is consistent with a global analysis (Belmaker et al., 2012). However, the strength of the richness-specialization relationship has strongly decreased strongly over the past 50 years, indicating that species richness has become increasingly decoupled from community-level specialization. In this study, land use and climate change are shown to induce a loss of contribution of community-level specialization to species richness, while species richness appears increasingly linked to local environmental conditions and their changes over time.

The weakening of the richness-specialization relationship is supported by different analyses in the present study, which all provide complementary perspectives on it. The dedicated analysis (step 1) informs about the trend of relationship itself. We then tested how the loss of the richness-specialization relationship is translated into changes in the spatial covariation of richness and specialization over time (step 2 and 3). Finally, the STATICO analysis (step 4) links the weakening of the richness-specialization relationship to its environmental drivers.

The loss of the richness-specialization relationship over time is consistent with the trend toward biotic homogenization, i.e. the local extirpation of specialists and/or gain in generalists, widely supported in the literature (Clavel, Julliard, & Devictor, 2011; Davey et al., 2012; McKinney & Lockwood, 1999; Newbold et al., 2018). Our results, which show that species richness increased over the 50 years of the study, are also consistent with previous findings demonstrating that local richness is not systematically decreasing over time (Dornelas et al., 2014; Hillebrand et al., 2017). Looking simultaneously at species richness and specialization trends, the present study thus confirms that changes in community composition, sometimes illustrated with species turnover, are a dominant response of communities to global changes (Dornelas et al., 2014; Hillebrand et al., 2017; Larsen, Chase, Durance, & Ormerod, 2018). This joined analysis indeed suggests that species specialization is an important driver of these compositional changes, but not always leading to the local extirpation of specialists.

Hence, beyond the general agreement between the weakening of the richness-specialization relationship and biotic homogenization trends, our results depict a more complex situation where biotic homogenization is a strong but not the only response of communities to global changes. This complex situation is captured by the first axis of the STATICO analysis, along which rich and low-specialized communities are getting richer and less specialized over time (biotic homogenization) while poor but highly specialized communities are getting more specialized and poor over time. This second trend, also visible in the decreasing slope of the second part of the piecewise regression (step 2), was observed in harsh environments free of human land use, where climate change induced a rise of harshness over time (mountainous areas). It is likely that such trend arises from low human impact on land that preserves specialists combined with rising climate harshness that tends to decrease niche space, locally selecting adapted (specialized) species but filtering generalists out (Attum, Eason, Cobbs, & Baha El Din, 2006). Because this specialization/species loss trend is not linked to a loss in specialization but to a niche reduction, it does not impact the richness-specialization relationship, i.e. the contribution of specialization to richness. However, it modifies the spatial covariation pattern of richness and specialization and therefore can influence our perception of the relationship between richness and specialization from a purely spatial point of view. Importantly, the STATICO analysis captures these two trends on the same axis (axis 1). First, this means that the “homogenization” and the “specialization through niche contraction” trends are tightly negatively correlated in the studied communities. Second, it also means that the environmental variables contributing to one trend also inversely contribute to the other, and that we cannot differentiate the drivers of both trends with our analyses.

Generally, our findings provide further support for specialization as a key explanation for macroecological diversity patterns, and thus support the idea that specialization, emerging from long-term stability and maintained by environmental stability is, or at least has been, an important driver of species richness (Currie et al., 2004; Gaston, 2000; MacArthur, 1972). In this theoretical context, the weakening of richness-specialization relationship as a macro-ecological pattern suggests that beyond the loss of specialization, high frequency and intensity of land use and climate changes on long time spans may impede diversification (Ponge, 2013).

### The drivers of richness and specialization covariation spatial pattern

Our study demonstrates that the positive richness-specialization relationship for US breeding birds over the past 50 years is spatially hidden by environmental and historical conditions, driving richness and community-level specialization. The spatial variation of those environmental variables translates into a nonlinear covariation pattern of richness and specialization, showing maximal richness at low to medium levels of specialization, a pattern also observed in Europe (Davey et al., 2012). Climate, land use, topographic and historical variables are all found to contribute to this nonlinear pattern.

Together with previous studies, our results tend to show that the more specialized communities are found in areas that experienced low climate velocity in the past and are still relatively preserved from human activities (Araújo et al., 2008; Dalsgaard et al., 2011; Sandel et al., 2011). These communities, forming the second part of the nonlinear covariation pattern, are found in harsh climate conditions of mountainous areas. Among those communities, the ones which are slightly used (rangelands) are less rich and specialized than the unused ones.

On the other hand, the richer communities are also found to be low-specialized. These rich communities are found in areas which experienced high climate velocity in the past, but are also intensively used by humans and beneficiate from a mild climate, favouring high levels of productivity. These results tend to indicate that the potential specialization level of those communities could be limited, due to high climate velocity in the past (Barratt et al., 2018; Dalsgaard et al., 2011; Sandel et al., 2011) and that this potential has been largely eroded by the intensive use of lands by humans (Clavel et al., 2011; Devictor et al., 2008). The human activities were already strongly driving the pattern during the first decade of the study, suggesting that the pattern has been shaped and altered by human activities for decades and probably centuries. Those impacts mainly concerned the most productive and accessible areas of the east, where the largest and prolonged effects of human development are situated in (Ellis et al., 2013; Sandom et al., 2014). The intensive use of land also drives a pattern of both poor and low-specialized communities, mainly represented in the less productive lands but also in areas totally dedicated to cropping, where intensified agriculture probably drastically lower available resources, leading to the observed richness and specialization decline. Intensive cropping drives the first part of the nonlinear covariation pattern, where the strong richness and slight specialization increases are linked to a decrease from high to medium cropping intensity, in term of land proportion and irrigation.

Thus, the spatial covariation pattern of richness and specialization for breeding birds in the US does not materialize along a latitudinal gradient, but rather follows multiple topographic, climate and land use gradients, all of which determine available productivity and environmental stability. This observation demonstrates the importance of directly studying the underlying biophysical gradients of the relationship instead of their latitudinal approximation to understand biodiversity patterns and their response (Gaston, 2000).

### Drivers of a weakening richness-specialization relationship through time

We show that the richness-specialization relationship weakened during the past 50 years in response to climate change, which confirms the driving role of climate change in community turnover previously observed in North-American bird communities (Barnagaud, Kissling, et al., 2017; Princé & Zuckerberg, 2015). The coarse spatial and thematic resolution of the land cover data prevented definitive conclusions on the importance of land use and land use intensity in weakening the richness-specialization relationship. Higher spatial and thematic resolution land use data would greatly improve our understanding of recent biodiversity trends.

We identified three main groups of variables that drove temporal change in the richness-specialization relationship and in the spatial covariation of richness and specialization. The first two groups contributed to a weakening of the relationship, while the third strengthened the relationship, which may explain the stabilisation of the strength of the relationship since the year 1990 (Figure 4).

(i) Increasing inter-annual temperature variation had the strongest negative impact on specialization and positive impact on richness (Gilchrist, 1995). Temperature effects were mainly concentrated in the East and North-East. Interestingly, our results show that the more specialized areas of the US (Rocky Mountains), have been relatively preserved from the increase in climate instability during the 50 years of the study, preserving specialized species.
(ii) The changes in winter and breeding season precipitation also altered the richness-specialization relationship. These changes mostly reinforced initial local conditions, increasing resource availability and richness in the already productive but heavily used areas of the East, while decreasing winter precipitation in the West. As a consequence, specialization increased while richness decreased in the West, and the opposite trend was observed in the East. In western communities, this suggests a loss of generalist species following increased ecological stress, which has also been documented elsewhere (Attum et al., 2006; Ponge, 2013).
(ii) A trend of rising naturalness was observed in points corresponding to the top left quarter of the STATICO analysis (Central Plains heavily used for agriculture; Figure 6a and 6c, Figure 7), which reduced the pressure on those communities and was beneficial for both richness and specialization. The way we computed the naturalness indicator suggests that this rise of naturalness corresponded to a decrease of croplands, suggesting a land abandonment process beneficial for late-successional species (Laiolo, Dondero, Ciliento, Animale, & Albertina, 2004).

## Conclusion

This study, for the first time, shows a decline in the strength of the richness-specialization relationship across recent decades. Importantly, we can link the weakening of this theoretically predicted macroecological pattern to recent climate and land use changes. These results show that human-driven land use and climate changes over are modifying macroecological patterns, changing the order of importance of processes that drive global diversity patterns. Thus, we show that community composition is increasingly affected by human-driven short term environmental instability and land use, which are now likely over-riding deep evolutionary patterns linked to the origination and long-term maintenance of species diversity. In addition to supporting previous finding on biotic homogenization, the joint analysis of richness and specialization proves also to be pertinent to capture other community response to global change, as niche contraction with specialization maintain as observed in this study.

## Acknowledgements

The authors wish to acknowledge the North American Breeding Bird Survey and all the volunteers that have been gathering data and organizing this great data base since more than 50 years. This work has been funded by the project BACI of the European Union H2020 Research and Innovation Program, under the number 640176. JCS considers this work a contribution to his VILLUM Investigator project “Biodiversity Dynamics in a Changing World” funded by VILLUM FONDEN (grant 16549).

## Data accessibility Statement

All data are directly downloadable online:

- BBS data: https://www.pwrc.usgs.gov/bbs/
- Globcover data: http://due.esrin.esa.int/page_globcover.php
- Biome data: https://dds.cr.usgs.gov/srtm/version2_1/SRTM30/w020n40/
- SRTM data: https://dds.cr.usgs.gov/srtm/version2_1/SRTM30/w020n40/
- Worldclim data: http://worldclim.org/version2
- HYDE data: http://themasites.pbl.nl/tridion/en/themasites/hyde/download/index-2.html
- NPPO (Haberl et al., 2007): https://www.aau.at/en/social-ecology/data-download/
- Climate velocity (Sandel et al., 2011): https://datadryad.org//resource/doi:10.5061/dryad.b13j1

## Appendix S1: The North American Breeding Bird Survey

The observations are made by volunteers who record bird abundances for three minutes within a radius of 400m along 50 fixed count points along a route. Count points are separated by at least 800 m. Observations are made during the peak of the breeding season, mainly in June. They begin 0.5 hour before local sunrise and last around 4 or 4.5 hours. Detail information is available and downloadable for the NABBS website (https://www.pwrc.usgs.gov/bbs/).

## Appendix S2: Simplification of the nomenclature of the Globcover data base used for estimating the specialization

The table below shows the Globcover nomenclature simplification used in the study to compute species habitat specialization. We excluded the points located in land cover classes that were not clearly dominated by one land cover (less than 50% of a unique identified habitat, i.e. GLC classes 30 and 110, where the main habitat can be different natural habitats). In grey are indicated the land cover classes that were not included because of the reason previously mentioned or because they concerned a too small number of count point data (<= 25 count points).

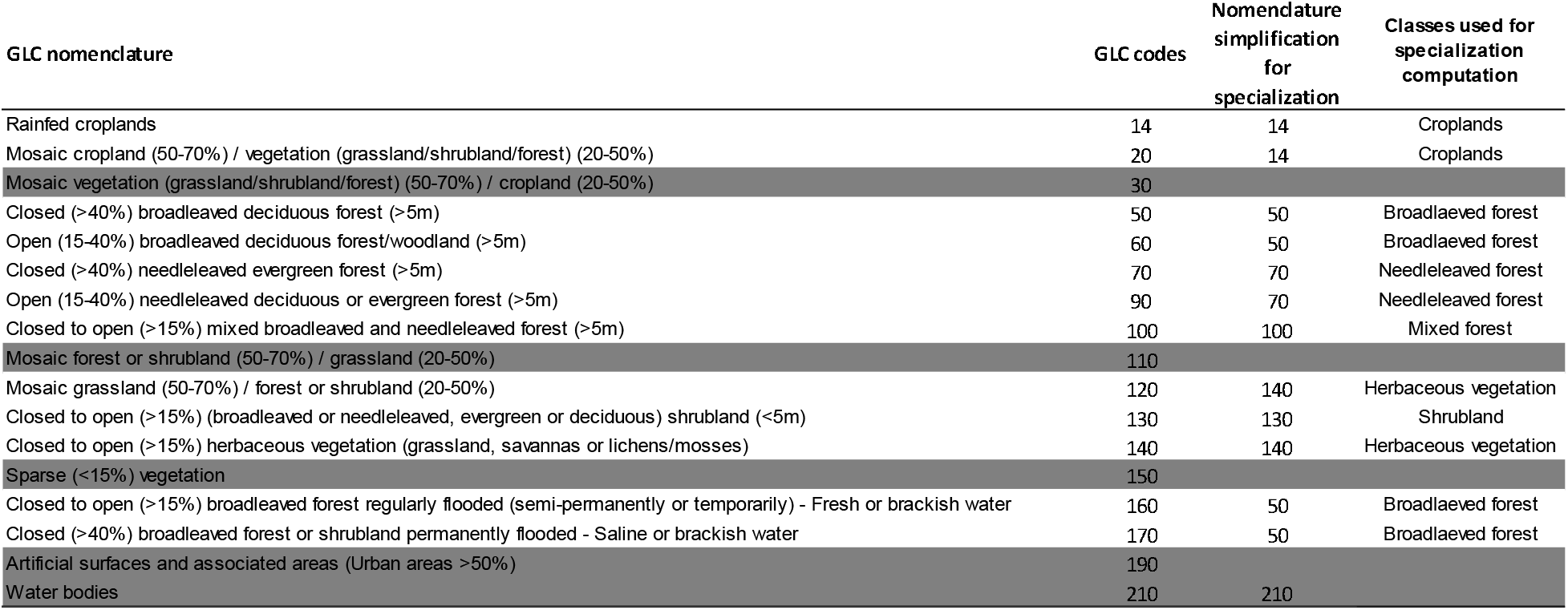

## Appendix S3: Structure of the different models used in the paper and associated R packages

| Temporal structure | Type | R Package | Shape of the Richness-Specialization relationship | Structure |
| --- | --- | --- | --- | --- |
| A unique model for all years | GLMM | glmmPQL | Linear | $Ric \sim SPE + year + SPE * year + 1 \mid id \ route$ |
| A model per year and by 11-years decade (three decades) | GAM | mgcv | Linear<br>Piecewise 1 (one breakpoint K)<br>Piecewise 2 (two breakpoints K1 and K2) | $Ric \sim SPE + s(X,Y)$<br>$Ric \sim SPE(0:K) + SPE(K:max(SPE)) + s(X,Y)$<br>$Ric \sim SPE(0:K1) + SPE(K1:K2) + SPE(K2:max(SPE)) + s(X,Y)$ |

## Appendix S4: Name, source and data transformation of the environmental variables used in the STATICO analysis

The data were extracted from buffers of 9500 meters around the routes. This radius corresponded to the resolution of our coarser variables (HYDE data). Climate data were available for each year of the decades. To characterize the climate within the buffers, we computed the average value of all years per pixel located in the buffer. From there, we quantified the average climate of the buffer as the mean. Then, as both spatial heterogeneity and temporal variations expected to impact species, we computed indicators for these two dimensions of change. For spatial heterogeneity, we calculated the standard deviation values between the pixels for all climatic variables (minimum and mean temperature and precipitation in winter and breeding season). For temporal stability, we focused on the climate variables of the breeding season, and first measured the mean yearly values within each buffer and then computed the standard deviation between years within the decade. We only had one year for each decade of land use data, and thus couldn’t compute indicators of temporal stability for land use.

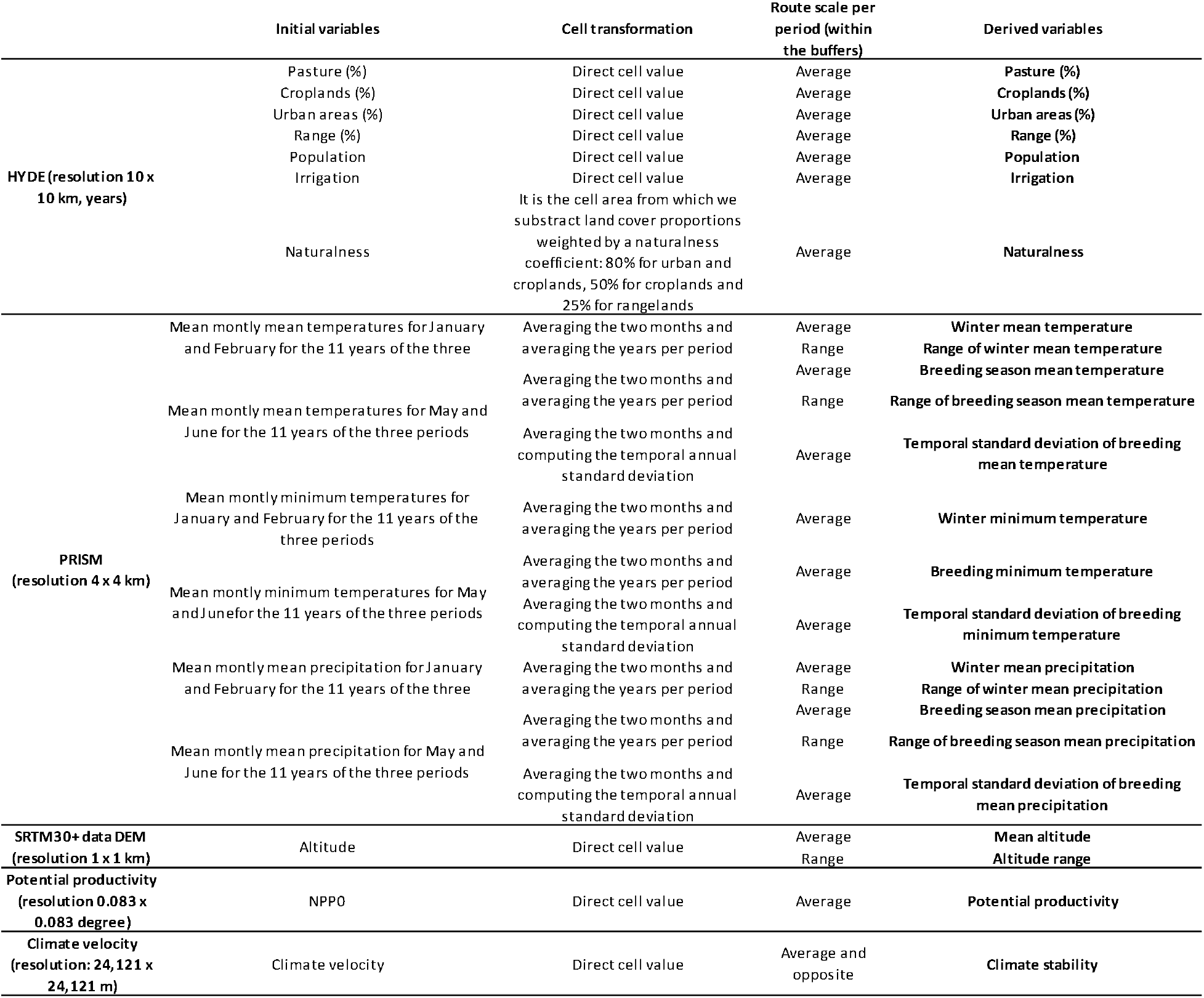

## Appendix S5: The STATICO analysis

In a first step, the STATICO method runs a co-inertia analysis between the response variables (species richness and community specialization) and the explanatory (environmental) variables for each decade (i.e. three decades in the present study). Co-inertia analysis is a multivariate method for coupling two tables, estimating the concordance between two tables based on the co-inertia criterion as measure of co-structure of the two tables. In our case it extracts the costructure between the table of richness and specialization on one side and the table of environmental variables on the other, and does it independently for the three decades. In a second step, the STATICO method runs a Partial Triadic Analysis (PTA). The PTA is a multivariate analysis adapted to study the variability of spatial or temporal structure between data tables with similar structure but varying in space or time. It also extracts a co-structure between the input tables (called the compromise), here the three co-structure tables from the three coinertia analyses. This compromise is contained into a multivariate space called the compromise space, from which the results are interpreted.

## Appendix S6: Results of the two breakpoints piecewise for the three decades

The figure shows the relationship between richness and specialization as obtained with the two breakpoints piecewise regression on the averaged data per decade. The standard errors are represented in shaded colors.

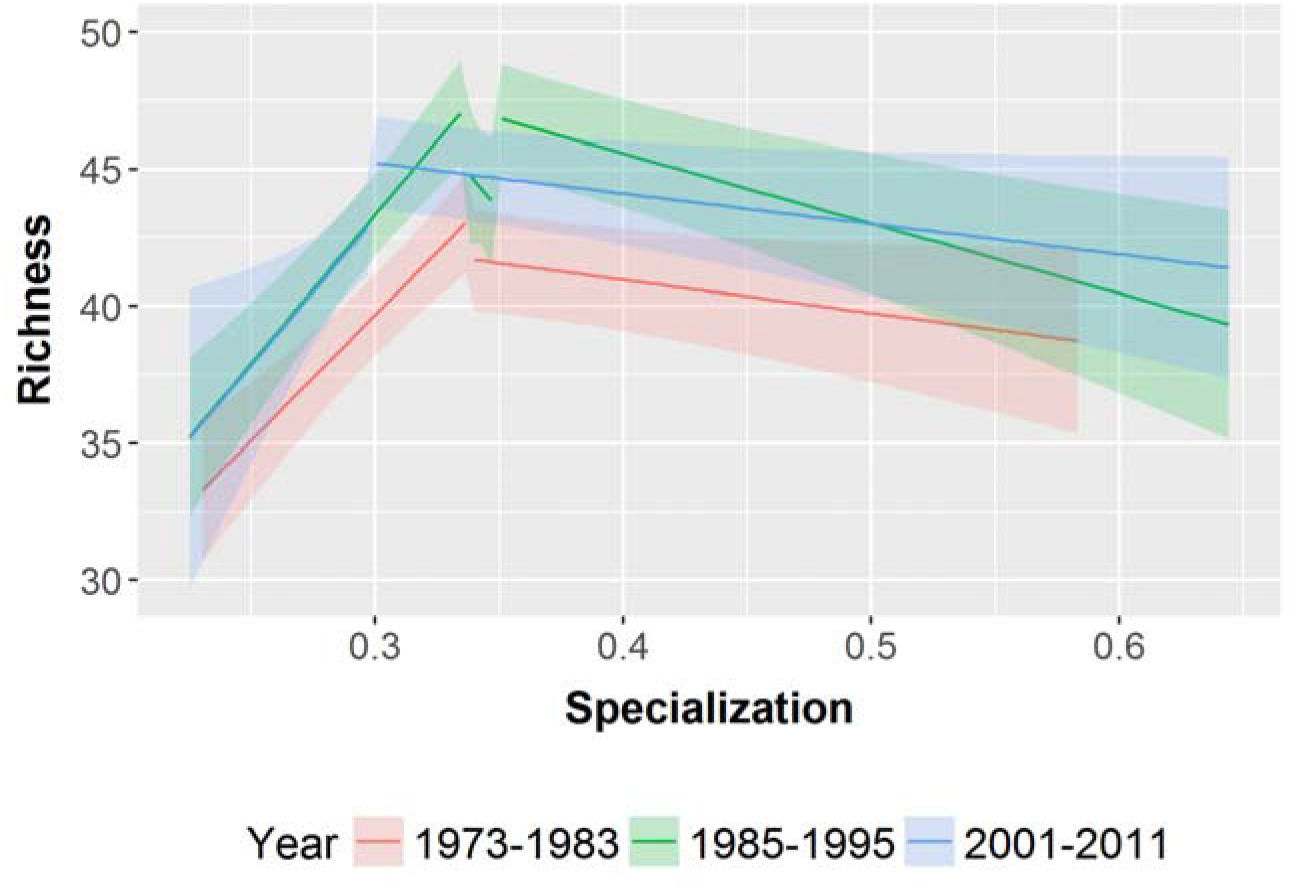

## Appendix S7: Complete interpretation of the results on the temporal dynamics of the STATICO analysis

### Changes in axis 1

The negative relationship between richness and specialization carried by the first axis became stronger in time, as richness and specialization moved away from each other along the axis. The reinforcement of the negative relationship was linked to change in temperature and precipitation regimes.

The importance of the inter-annual variations in minimum temperature increased in time while climate change strengthened the pre-existing spatial patterns (Fig 6c). From being linked to higher specialization values in the first decade (negative part of axis 1), the Minimum temperature (winter) shifted toward being linked to higher richness values during the last decade (positive part of axis 1). This shift occurred as the the Minimum temperature (winter) decreased in the negative part of axis 1 and increased in the positive part of axis 1, actually reversing the initial pattern. The importance of mean precipitation during the breeding season increased in time as climate change induced a reinforcement of the initial conditions, i.e. more precipitation for the wetter areas (positive part of axis 1) and less precipitation for the drier routes (negative part of axis 1) (Figure 6c).

## Changes in axis 2

Meanwhile, the slight positive relationship between richness and specialization carried by the second axis became weaker as the correlation of the two variables with the axis decreased (Figure 6b). These changes were related to changes in climate and land-use intensification. The importance of both crops and naturalness decreased in time as cropping globally lowered and naturalness increased in the intensified warm and dry regions (poor and generalist communities) (Figure 4c). These changes co-occurred with decreasing importance of precipitation in winter and of the spatial range of precipitation in winter, as initial patterns weakened with recent climate change (Figure 6 b and c).

